# Efficient, Few-shot Directed Evolution with Energy Rank Alignment

**DOI:** 10.64898/2026.02.03.703561

**Authors:** Sebastian Ibarraran, Shriram Chennakesavalu, Frank Hu, Grant M. Rotskoff

**Affiliations:** Department of Chemistry, Stanford University, Stanford CA 94305; Institute for Computational and Mathematical Engineering, Stanford University, Stanford CA 94305

## Abstract

Directed evolution is a powerful and widely used technique for protein engineering, and reducing the cost of iterated experimental observations has become a major priority for practitioners. A number of recent efforts to use machine-learning-based predictors to improve sequence selection have led to remarkable improvements in efficiency, but the sparse data at each experimental iteration restricts these approaches to extremely simple models. Adapting large-scale pre-trained protein language models using experimental data offers an alternative that we show productively leverages the strong inductive biases of the natural distribution of protein sequences to navigate high-dimensional, combinatorially large fitness landscapes. Our approach uses a general-purpose “post-training” algorithm grounded in statistical physics that employs quantitative experimental rankings to directly produce a sampler for diverse, high fitness sequences with fewer data points than competing methods. The resulting adapted protein language model can itself be studied and interpreted, shedding further light on the biophysical characteristics of highly fit sequences and their properties.

## Introduction

The advent of highly accurate structure prediction and sequence design tools has refocused protein engineering on structural features that strongly correlate with desired function [1, 45, 17, 8], but these methods do not directly leverage relevant experimental data when generating predictions. Experimental design via directed evolution [44] (DE) stands somewhat in contrast to this trend; in DE, an experimental observable guides the sequential perturbation of a parent sequence. There is a productive and relatively unexplored interface between these two approaches, and a variety of recent methods have sought to augment DE with data-driven machine learning [47, 52]. Choosing candidate sequences in DE is complicated by a combinatorially large fitness landscape and epistatic effects that are challenging to anticipate *a priori* [41, 21]. Integrating DE workflows with the powerful inductive biases learned by large-scale pre-trained statistical models for sequences, such as protein language models [25, 15], offers a compelling route to navigating rugged, high-dimensional fitness landscapes.

The number of measurements used in standard DE protocols remains modest and, as a result, previous efforts to integrate machine learning into DE have relied on simple model architectures suitable for small quantities of data [35, 36, 33, 2]. While these strategies have demonstrated an improvement over conventional DE, the models lack the strong inductive bias of large-scale pre-trained models, which limits their ability to capture the complex correlations in protein sequence that underlie fitness. To mitigate this loss of representation complexity, predictors have relied on orthogonal measures of protein fitness, as the models themselves do not uniformly correlate well with fitness [27, 14, 11]. More sophisticated models, however, require significantly more data than is typically available in a single experimental observation step [5].

Protein language models (PLMs) trained on billions of protein sequences [25, 15, 4, 13] could mitigate this lack of data by providing a strong evolutionary inductive bias. PLMs are built using transformer architectures, which directly parameterize a probability distribution over the sequences of naturally occurring proteins. Furthermore, it is well-known that sequence statistical likelihoods strongly correlate with thermodynamic stability [23, 29], which has been leveraged to predict relative sequence fitnesses for protein-protein interactions [9]. PLM likelihoods have also been used as a constraint to identify high fitness, both with and without structure conditioning [16, 39]. However, the robustness of the assumption that the sequence likelihoods and experimental activities are strongly correlated is not appropriate in all protein optimization settings, especially those far from natural function. For experimentally guided design tasks, folding stability leads to a probability distribution that is too diffuse to productively guide sampling of candidate sequences.

Adapting PLMs with small amounts of experimental data closely resembles a widely studied problem in the machine learning literature known as “post-training”, in which a pre-trained model is optimized to yield a desired distributional shift. While many post-training methods exist [38, 30], preference optimization [32, 46] is uniquely well-suited to the comparative task of ranking sequences using quantitative experimental outputs. Preference optimization seeks to adapt the model probabilities to impose a relative ranking among outputs that reflects an observed ranking; we recently developed a method called energy rank alignment (ERA) that ensures that relative probabilities among quantitatively ordered samples are preserved [6, 7]. ERA is an explicit gradient-based algorithm which can be implemented efficiently to post-train even very large PLMs.

Here we show that predicted sequences using post-trained PLMs efficiently optimize fitness across a variety of tasks: maximizing enzymatic activity, antibiotic resistance, and protein-protein binding. ERA-adapted protein language models sample sequences that identify the highest fitness sequences faster and more robustly than alternative techniques that combine simple models and active learning, including EVOLVEPro, Active Learning-assisted Directed Evolution, Machine Learning Directed Evolution, and Direct Preference Optimization [20, 51, 50, 48, 32]. Interestingly, augmenting pre-trained PLMs with 3D structural information or thermodynamic stability data does not outperform a predictor that purely relies on protein sequence. We exhaustively benchmark our approach using previously collected combinatorially complete experimental data sets with notable epistatic effects. We consistently see that ERA shifts the sequence distribution towards high activity but importantly maintains diversity even after post-training optimization.

One marked advantage of the approach we pursue is the increasingly formalized science of interpretability for language model architectures. By analyzing the distributions over amino acids for the adapted generators, we can compare highly fit sequence distributions to the native baseline. This in turn enables analysis of the physiochemical changes required to maximize performance on a given protein engineering task. While the relative computational effort required to optimize large-scale machine learning models like state-of-the-art PLMs is high compared to methods that use simple architectures, the conceptual and practical advantages make a strong case for expending the additional effort, given the even greater experimental costs.

## 1 Approach

### 1.1 Sampling mutants from the PLM

We aim to improve the efficiency of directed evolution campaigns where a fixed set of residues are chosen for mutation by practitioners and only *N* ≈ 100 are available at each iteration. Focusing on this setting enables rapid assessment of the quality of sequences generated by a PLM prior, as a number of combinatorially complete fitness landscapes have been published [31, 49, 26, 22]. We intentionally do not study higher throughput protein optimization methods such as phage-assisted continuous evolution (PACE) [12], because the mutations are made at random by an error-prone polymerase and these methods require building intricate reporter circuits [28].

Sampling mutant sequences requires conditionally evaluating amino acid likelihoods at residue positions of interest. Bidirectional transformers, also called masked language models [10], are well suited to this task because we can mask positions of interest and resample them. Throughout, we employ ESM3-1.4B [15], a state-of-the-art PLM that uses a bidirectional transformer architecture. Furthermore, the multi-modal nature of the model allows for additional conditioning beyond amino acid sequences, such as backbone coordinates, secondary structure elements, and even functional annotations. Given that the positions of interest for this study were largely chosen according to some structural intuition, we hypothesized that incorporating the coordinates of known crystal structures into the sequence design process would enhance predictive power.

To sample sequences from ESM3-1.4B, we mask the residue indices of interest within the protein target. These positions were then unmasked *simultaneously*, rather than permuting all possible unmasking orders between multiple sites. This enables us to treat the unmasking of token *x_i_, i* = 1*, …, n* at each of the *n* positions as statistically independent events and compute generation pseudo-likelihoods from the ESM3-1.4B policy *π*_ESM3_ using this unmasking scheme as:

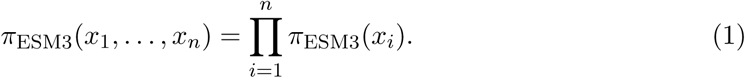

Exact likelihood calculations for arbitrary unmasking order in masked language models remains a topic of significant research interest [37, 42].

### 1.2 Adaptation with Energy Rank Alignment

To fine-tune ESM3-1.4B in a manner that leverages experimental fitness readouts of generated sequences, we choose to apply preference optimization due to its contrastive signal. With a batch of *N* sequences sampled from *π*_ESM3_, where *N* ≪ 1000 (never exceeds 384 in this work), we construct the full dataset of *^N^*^(*N*−1)^ preference pairs labeled with their experimental readouts, as illustrated in Fig. 1. Our goal is to minimize the divergence between the parametric preference distribution *p_θ_*:

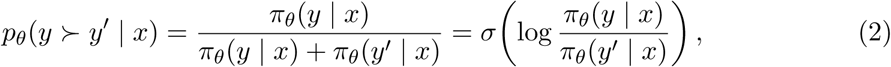

and the entropy regularized preference distribution *p_γ_*:

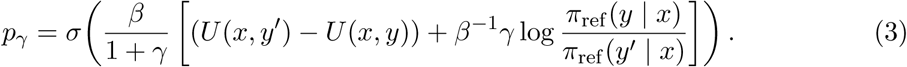

Here *β* tunes the entropy of the Gibbs-Boltzmann target distribution, *γ* is a regularization hyperparameter, and *U* (*x, y*) is an energy function (negative reward) to be minimized. The notation *y* ≻ *y*^′^ indicates that *y* is preferred to *y*^′^. We optimize the following KL-divergence, the minimizer of which coincides with (3),

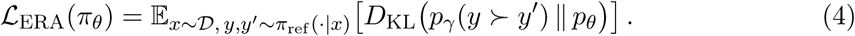

The choice of ERA as a fine-tuning algorithm for ESM3-1.4B provides several benefits compared to other methods. Continued training of *π*_ESM3_ with unlabeled sequences through supervised fine-tuning (SFT) would rely upon only selecting high-fitness sequences, which are exceedingly rare in more challenging protein fitness landscapes. The contrastive nature of a preference optimization approach leverages useful information from low-reward completions that would be otherwise discarded in an SFT approach that only uses high-reward sequences. Another potential strategy is reinforcement learning (RL), which would leverage the experimental readouts as a reward. The simple, gradient-based objective posed by ERA is computationally efficient compared to many RL algorithms such as proximal policy optimization (PPO), which operate within an actor-critic framework. Furthermore, the ERA objective also ensures sample diversity, which is a pitfall of many RL methods that can be prone to reward hacking and mode collapse. Previous work [6, 7] has shown that ERA outperforms standard preference optimization and RL techniques for continuous reward signals.

**Figure 1:**
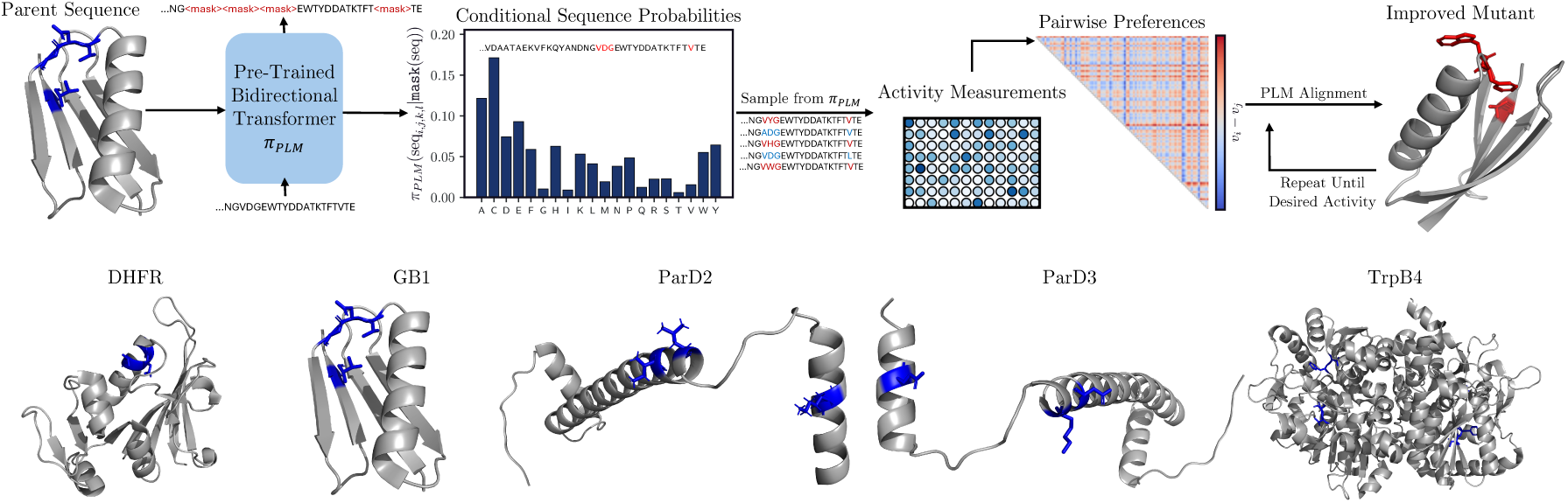
Preference alignment of a protein language model for efficient protein optimization in a directed evolution campaign. Using the conditional sequence distribution over residues, an initial set of protein sequences is sampled and evaluated. These sequences are compiled into a dataset of pairwise preferences labeled by their experimental fitness. This dataset is used to fine-tune the model with the energy rank alignment objective, targeting high-fitness sequences with tuneable entropy and controllable regularization. This process can be repeated iteratively in an active learning framework until some desired level of sequence fitness is achieved.

Our reward function is the experimental fitness readout associated with sequence *y*, i.e., *U* (*x, y*) = − log(*f_y_*), where *f_y_* is fitness on the unit interval after being scaled to the maximum fitness in the landscape. In each of our experiments, the model was trained using 4 for 25 epochs, which ensured that the training loss had converged. All parameters of ESM3-1.4B were trainable throughout this process, though techniques such as low-rank adaptation (LoRA) [18] could be employed to reduce the memory demands of training. We chose not to explore parameter-efficient variants of ERA in this work, but recent research has shown that LoRA can yield results in line with full fine-tuning [40]. With this aligned model, we sampled *N* sequences from the updated *π*_ESM3_ and their experimental fitness readouts were obtained. Duplicate sequences and sequences explored in previous rounds are excluded. Furthermore, we recompute the likelihoods of sequences sampled in previous rounds according to 1 using the updated *π*_ESM3_ as the policy. At each round of optimization, a new pairwise preference dataset is constructed with all available sequences and their updated likelihoods, and the process is repeated. In order to clearly compare with existing methods, the iterative loop was only repeated for four rounds in this study, but this process could be repeated until any threshold for desired activity is achieved.

## 2 Results

### 2.1 Comparison to Related Methods

We compare ERA to baselines for ML-driven DE, which vary both in choice of architectures and sequence representation. Machine learning directed evolution (MLDE) [50, 48] trains XGBoost and ridge regression models on either one-hot representations or ESM2-650M embeddings to predict sequence activities based on those that have been allowed by the sampling budget. The activities of all sequences in the landscape are then predicted, and the top *N* are selected according to these predictions. Active Learning-assisted Directed Evolution (ALDE) [51] is an extension of MLDE that uses boosting and deep neural network ensembles as the underlying architecture. Rather than training for a single round like MLDE, ALDE instead uses a greedy acquisition function to select sequences for evaluation and further training in an active learning framework. EVOLVEpro [20] can be viewed as a variant of ALDE that uses a random forest as the architecture for the activity prediction model and the top *N* sequences are naively sampled as in MLDE. As the original EVOLVE-pro study did not involve the same fitness landscapes that are investigated here, we took the opportunity to compare to EVOLVEpro trained on both ESM2-650M and ESM3-1.4B embeddings to assess the differences in performance between the two models. Lastly, we also compare ERA to Direct Preference Optimization (DPO) [32] on ESM3-1.4B, another preference optimization algorithm that differs from ERA in that it only uses rank ordering, not numerical fitness values. The comparison to DPO highlights the difference in performance afforded by labeling the magnitude of the pairwise preferences and the tunability of fluctuations around the optimal reward in the target distribution of ERA.

In order to benchmark the suitability of ERA for DE campaigns, we used previously collected experimental DE datasets that were nearly combinatorially complete. Li et al. collected a set of diverse protein activity landscapes across five different protein targets which were well-suited for this task [24]. These data feature a range of protein sizes, families, and activities, ranging from antibody binding to enzymatic efficiency. Three of the datasets—dihydrofolate reductase (DHFR) trimethoprim binding [31] and the ParD2/3-ParE2/3 bacterial antitoxin-toxin binding landscapes [26] —involve mutating three positions of interest combinatorially for a total of 8,000 possible sequences. Two other datasets—the immunoglobulin binding of the B1 domain of protein G (GB1) [49] and the tryptophan production rate of the *β*-subunit of tryptophan synthase (TrpB4) [22]—involve mutating four positions, making for 160,000 possible sequences.

Across all landscapes, four rounds of training with 96 sequences each were conducted followed by a final 96 sequences, for a total of 480 samples. The only exception to this was MLDE, for which only a single round of training on 384 sequences was conducted followed by sampling of 96 sequences as in [24]. The sampling process across all methods and replicates allowed only for unique sequences that had not been previously trained on, except for DPO. Significant mode collapse in the DPO-trained ESM3-1.4B model required us to allow for repeated samples in order to make the benchmark tractable.

We compare these methods according to two metrics outlined in [24]. The first is the average maximum fitness achieved across multiple replicates of these simulated DE experiments. The second is the fraction of replicates that reach the highest activity sequence. These two metrics provide insight into how well each method is able to fully optimize a protein sequence and their ability to locate diverse, high-activity sequences within a large fitness landscape.

Table 1 and Table 2 demonstrate that ERA attains state-of-the-art performance across a variety of diverse combinatorial fitness landscapes. The superior performance across the board compared to MLDE demonstrates the advantages of an iterative, active learning-type approach where the reference policy is updated more frequently. The inclusion of continuous labels in the training process leads to clearly superior performance compared to DPO. The mode collapse experienced by ESM3-1.4B when trained with DPO was so severe that 9 of the 10 replicate experiments converged to the exact same local optimum for TrpB4: V183; F184I; V227M; S228G. The tuneable entropy and explicit treatment of the experimental labels for each sequence allowed ERA not only to sample diverse high-fitness sequences, but also to navigate out of local optima in rugged fitness landscapes.

**Table 1:** Average maximum fitness achieved across multiple replicate runs with standard errors included. All methods used 384 total samples for training, with ALDE, DPO, and ERA doing so over four rounds (96 per round). For MLDE and ALDE, 50 replicates were averaged and values were obtained from [24], whereas 10 replicates were averaged for ERA, DPO, and EVOLVEpro.

| Method | DHFR | GB1 | ParD2 | ParD3 | TrpB4 |
| --- | --- | --- | --- | --- | --- |
| MLDE | 0.96 | 0.67 | 0.99 | 0.9 | 0.78 |
| ALDE | 0.96 | 0.80 | 0.99 | 0.99 | 0.89 |
| EVOLVE<sub>pro</sub> (ESM2) | 0.95 | 0.81 | **1.0** | 0.99 | 0.88 |
| EVOLVE<sub>pro</sub> (ESM3) | **1.0** | 0.77 | **1.0** | 0.99 | 0.79 |
| DPO | **1.0** | 0.78 | **1.0** | 0.99 | 0.81 |
| ERA | **1.0 ± 0.00** | **0.83 ± 0.04** | **1.0 ± 0.00** | **1.0 ± 0.00** | **0.91 ± 0.02** |

**Table 2:** Fraction of replicate runs reaching the maximum fitness sequence in the landscape. All methods used 384 total samples for training, with ALDE, DPO, and ERA doing so over four rounds (96 per round). For MLDE and ALDE, 50 replicates were averaged and values were obtained from [24], whereas 10 replicates were averaged for ERA, DPO, and EVOLVEpro.

| Method | DHFR | GB1 | ParD2 | ParD3 | TrpB4 |
| --- | --- | --- | --- | --- | --- |
| MLDE | 0.60 | 0.06 | 0.98 | 0.12 | 0.02 |
| ALDE | 0.62 | 0.18 | 0.96 | 0.80 | **0.46** |
| EVOLVE<sub>pro</sub> (ESM2) | 0.00 | 0.10 | 0.80 | 0.70 | 0.40 |
| EVOLVE<sub>pro</sub> (ESM3) | 0.90 | 0.00 | 0.60 | 0.10 | 0.00 |
| DPO | **1.0** | 0.10 | 0.90 | 0.00 | 0.00 |
| ERA | **1.0** | **0.20** | **1.0** | **0.90** | 0.30 |

The comparison to EVOLVEpro was insightful due to the difference in its performance depending on whether ESM2-650M or ESM3-1.4B embeddings were used to train the random forest. For all landscapes except DHFR, the performance was notably superior when using the ESM2-650M embeddings. This raises compelling questions about whether the latent representation of proteins learned by ESM3-1.4B is most suitable for sequence-level optimization tasks. Here, we primarily focus on ESM3-1.4B due to its generative capabilities and the flexibility to condition on different modalities like 3D structure, but ERA is agnostic to what the reference policy is and aligning ESM2-650M would also be a reasonable approach.

Lastly, comparison to ALDE and the ESM2-650M version of EVOLVEpro does show ERA falling short in terms of full optimization of the TrpB4 sequence. Nevertheless, ERA marginally outperforms these methods in terms of the average maximum fitness achieved. This suggests that even in cases where ERA is not able to fully optimize the sequence in the allowed 480-sample budget, many of the other high-activity sequences are still recovered. In biological settings where *in vitro* to *in vivo* translation can be unreliable, this property of ERA can be advantageous to generate a larger number of high-quality samples and is consistent with previous observations about its behavior [7].

### 2.2 Assessing The Landscapes

After comparing ERA with other methods across a range of combinatorial fitness landscapes, we sought to assess the studied fitness landscapes to understand the differences in performance for ERA across them. To accomplish this, we use GraphFLA [19] to compute metrics related to the roughness and navigability of the landscapes. Table 3 compiles the local optima ratio (LOR), roughness-slope ratio (R-S ratio), neutrality index, and global optimum accessibility (GOA) of the five landscapes of interest. These metrics outline a clear hierarchy of difficulty in navigating the landscapes. The three-site landscapes—ParD2, ParD3, and DHFR—have a relatively low prevalence of local optima, possess accessible global optima, and have the lowest deviation from an additive model as quantified by the R-S ratio. The lower neutrality indices of ParD2 and ParD3 may seem like a potential obstacle to their evolvability, but this is largely avoided because the parent sequences for these landscapes are already close to the optimal activity. The four-site landscapes, GB1 and TrpB4, appear more challenging across most of these metrics in addition to simply consisting of a larger search space of sequences. The metrics associated with GB1 generally favor the navigability of that landscape compared to TrpB4, which may contribute to the superior performance of ERA for GB1 compared to TrpB4.

**Table 3:** Descriptive statistics of the roughness, neutrality, and navigability of combinatorial multi-site fitness landscapes. The computed quantities are the local optima ratio (LOR), roughness-slope ratio (R-S ratio), neutrality index, and global optimum accessibility (GOA). All quantities were obtained using landscapes from [24] and are computed using GraphFLA [19].

| **Landscape** | **LOR ↑** | **R-S ratio ↑** | **Neutrality Idx. ↓** | **GOA ↓** |
| --- | --- | --- | --- | --- |
| ParD2 | 0.0006 | 2.5681 | 0.0600 | 0.9995 |
| ParD3 | 0.0005 | 2.5889 | 0.0528 | 0.9996 |
| DHFR | 0.0004 | 5.8314 | 0.8274 | 0.9994 |
| GB1 | 0.0013 | 9.0128 | 0.8779 | 0.9976 |
| TrpB4 | 0.0050 | 29.6815 | 0.3430 | 0.9876 |

Table 4 expands on this analysis to include epistatic effects. Epistasis is a phenomenon where the effects of mutations at different sites are non-additive and is the primary challenge facing conventional techniques for sampling new sequences in DE campaigns. Specifically, we tabulate magnitude epistasis (non-additive effects where the sign stays the same), sign epistasis (the sign of one mutation effect flips in the presence of the other), and reciprocal sign epistasis (the sign of both mutation effects flips in the presence of the other). We observe similar trends as above: for instance, ParD2 and ParD3 have the most positive prevalence of magnitude epistasis, which should enhance evolvability. These fitness landscapes also have a considerably lower prevalence of sign and reciprocal sign epistasis than the others. While DHFR is comparably epistatic to the GB1 and TrpB4 landscapes, these effects are likely far easier to learn in a lower-dimensional sequence landscape than they are for the other two targets. The prevalence of strong epistatic effects across these landscapes is a challenge that a method like ERA, which promotes sample diversity and learns using rich latent representations of protein sequences, is well-suited to overcome, especially in comparison to greedier sampling schemes.

**Table 4:** Descriptive statistics of the epistatic effects present in combinatorial multi-site fitness landscapes. The prevalence of magnitude, sign, and reciprocal sign epistasis are tabulated. All quantities were obtained using landscapes from [24] and are computed using GraphFLA [19].

| **Landscape** | **Magnitude ↓** | **Sign ↑** | **Reciprocal sign ↑** |
| --- | --- | --- | --- |
| ParD2 | 0.5805 | 0.3069 | 0.1126 |
| ParD3 | 0.6522 | 0.2803 | 0.0676 |
| DHFR | 0.4013 | 0.3689 | 0.2298 |
| GB1 | 0.4349 | 0.3486 | 0.2165 |
| TrpB4 | 0.3598 | 0.3542 | 0.2860 |

### 2.3 Assessing Model Priors

In addition to examining the landscapes themselves, we also quantified the conditional distribution learned over sequences by ESM3-1.4B. Table 5 provides descriptive statistics of the *π*_ESM3_ distribution for the five landscapes as computed in 1. In addition to common metrics such as the entropy and Kullback-Leibler divergence of the distribution to a uniform distribution, we also quantify the probability mass concentrated in the five most likely sequences (Top-5 Coverage) and the prevalence of redundant sequences in a batch of 1000 samples (Redundancy). These metrics reflect similar trends to the epistatic effects from Table 5. ParD2, ParD3, and GB1 seem to have the *π*_ESM3_ distributions with the most spread, making them a less challenging prior to align the model away from. On the other hand, DHFR and TrpB4 have far more mode-seeking distributions according to these metrics that are more difficult to alter with fine-tuning. However, the key to ERA doing this successfully in this setting is the use of more sequences relative to the total size of the landscape than are available for TrpB4.

**Table 5:**
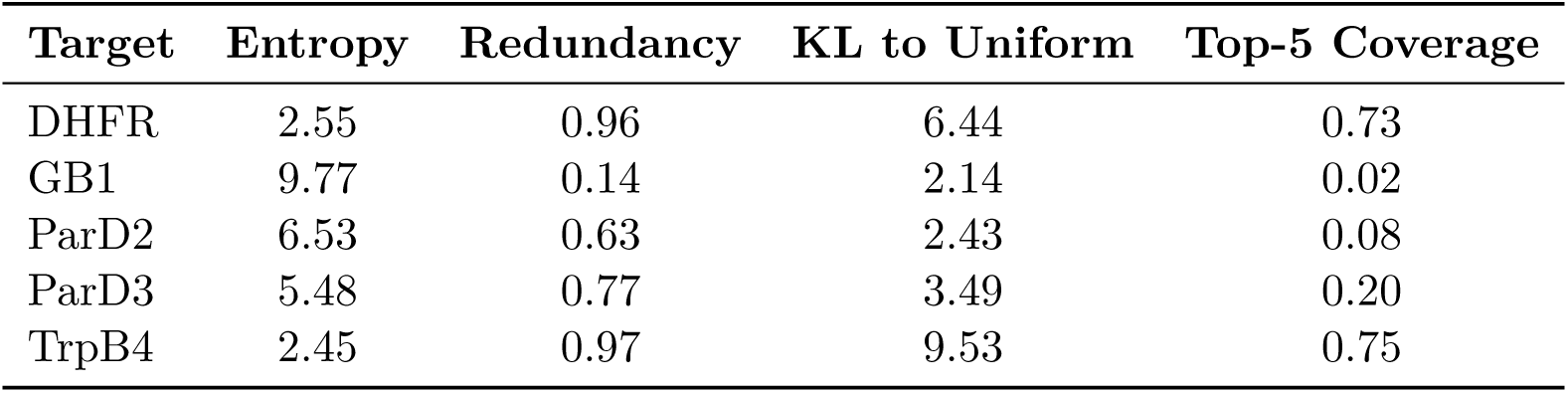
Descriptive statistics of the conditional distribution over sequences learned by ESM3-1.4B for five combinatorial multi-site protein fitness landscapes. These quantities - the entropy, sampling redundancy, KL to the uniform distribution, and probability mass covered in the top-5 sequences - quantify the extent of mode-seeking behavior in these distributions.

### 2.4 Effects of Alignment

Table 6 shows that unaligned log *π*_ESM3_ is largely uncorrelated with the experimental fitness values across the five landscapes. This relates to previous observations in the literature that the evolutionary fitness landscapes learned by PLMs are not necessarily zero-shot predictors of experimental activity, especially in settings where the activity being optimized is further from a natural function [20]. The goal of ERA is to use relatively small batches of experimental data to improve the agreement between these landscapes and make the PLM prior more useful for sampling sequences toward some desired functional optimization. For the targets that had a *π*_ESM3_ distribution with greater spread—namely GB1, ParD2, and ParD3—the correlation between log *π*_ESM3_ steadily improved with further rounds of alignment. On the other hand, the more mode-seeking log *π_ESM_*_3_ distributions of DHFR and TrpB4 experienced an initial dip in the correlation following the first round of alignment followed by an increase during subsequent iterations.

**Table 6:** Pearson correlations of sequence log *π*_ESM3_ with the corresponding experimental activity for five protein targets of interest. log *π*_ESM3_ were computed from a random iterative alignment experiment for that target.

| Target | Unaligned Corr. Coeff. | Aligned Corr. Coeff. (1 Round) | Aligned Corr. Coeff. (2 Rounds) | Aligned Corr. Coeff. (4 Rounds) |
| --- | --- | --- | --- | --- |
| DHFR | 0.547 | 0.470 | 0.635 | - |
| GB1 | 0.160 | 0.293 | 0.334 | 0.355 |
| ParD2 | 0.466 | 0.862 | - | - |
| ParD3 | 0.550 | 0.705 | 0.758 | 0.766 |
| TrpB4 | 0.361 | 0.309 | 0.370 | 0.334 |

Figure 2 visually demonstrates this change in correlation over several rounds of alignment. Fine-tuning ESM3-1.4B with ERA clearly suppresses the likelihoods of many inactive sequences and promotes the likelihoods of highly active sequences by several orders of magnitude. These correlations between log *π*_ESM3_ and the experimental activities are particularly impressive for the ParD2 and ParD3 landscapes, and the improvement for DHFR and GB1 is notable as well, despite the continued presence of some false-positives, i.e. sequences with high-likelihoods and low activities. Though the overall correlation for TrpB4 improves, the bimodal behavior of the log *π*_ESM3_ distribution in the aligned plot seems to suggest the presence of the local optima in the fitness landscape being located by training with ERA. Despite this, the inherent stochasticity of sampling directly from *π*_ESM3_ along with the sample diversity promoted by ERA still allows for the possibility of escaping these local optima when sampling sequences.

**Figure 2:**
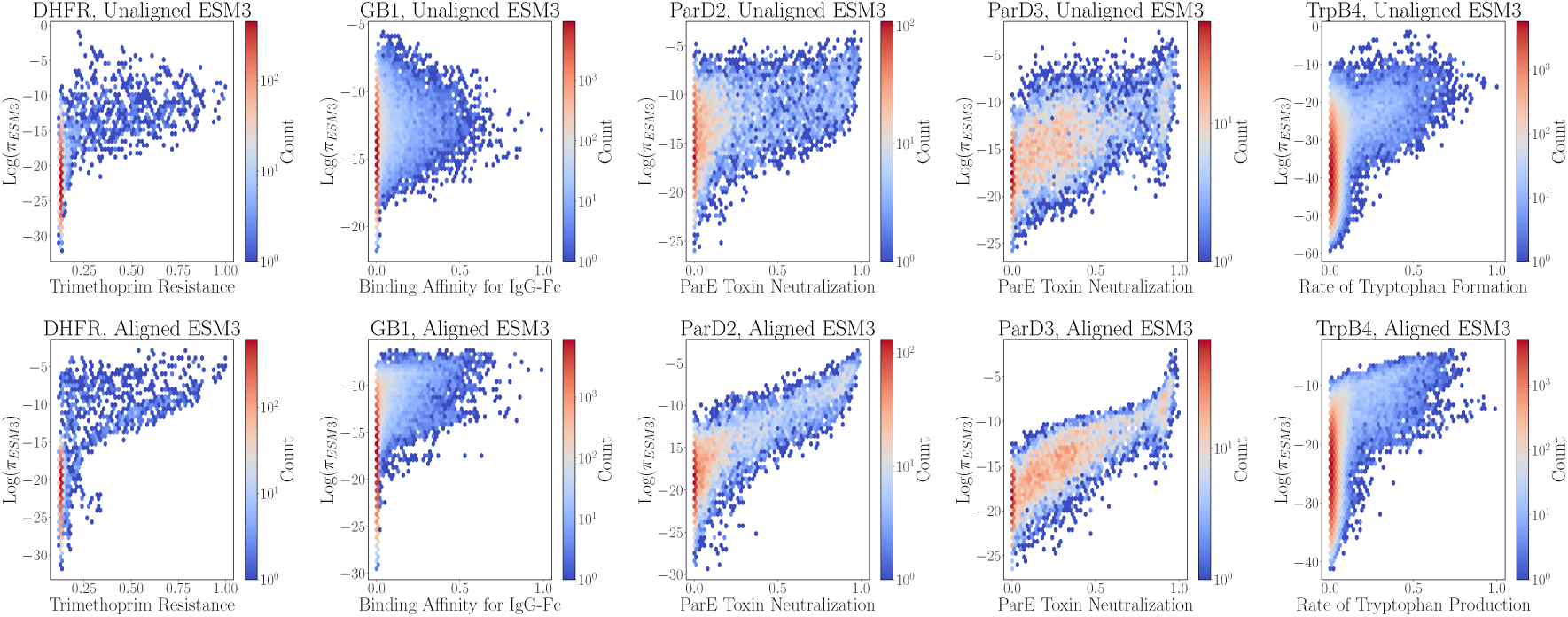
Evolution of the relationship between log *π*_ESM3_ and experimental activity from before (top row) to after (bottom row) alignment. The aligned models used to compute log *π*_ESM3_ in the bottom row are the final checkpoints in a random iterative alignment experiment.

The probability mass assigned to high-activity sequences under the post-ERA policy *π*_ESM3_ increases markedly. Figure 3 displays the fold-improvement (on a log-scale) of the probability mass assigned to sequences above a specified lower bound in activity after fine-tuning with ERA. More probability mass is assigned to high-activity sequences for every landscape, with the gains for DHFR, GB1, and TrpB4 appearing to be particularly substantial. Even for ParD2 and ParD3, whose fold-increases in probability mass look less impressive for their most active sequences, the relatively high probability mass assigned to those regions initially (0.25 above fitness of 0.9 for ParD2, 0.40 above fitness of 0.9 for ParD3) means these are still considerable absolute gains in probability mass for the high-activity region of sequence space. These distributional shifts are clearly reflected in the performance metrics from Tables 1 and 2, which demonstrate that the aligned policy is effectively able to sample sequences in these high-fitness regions.

**Figure 3:**
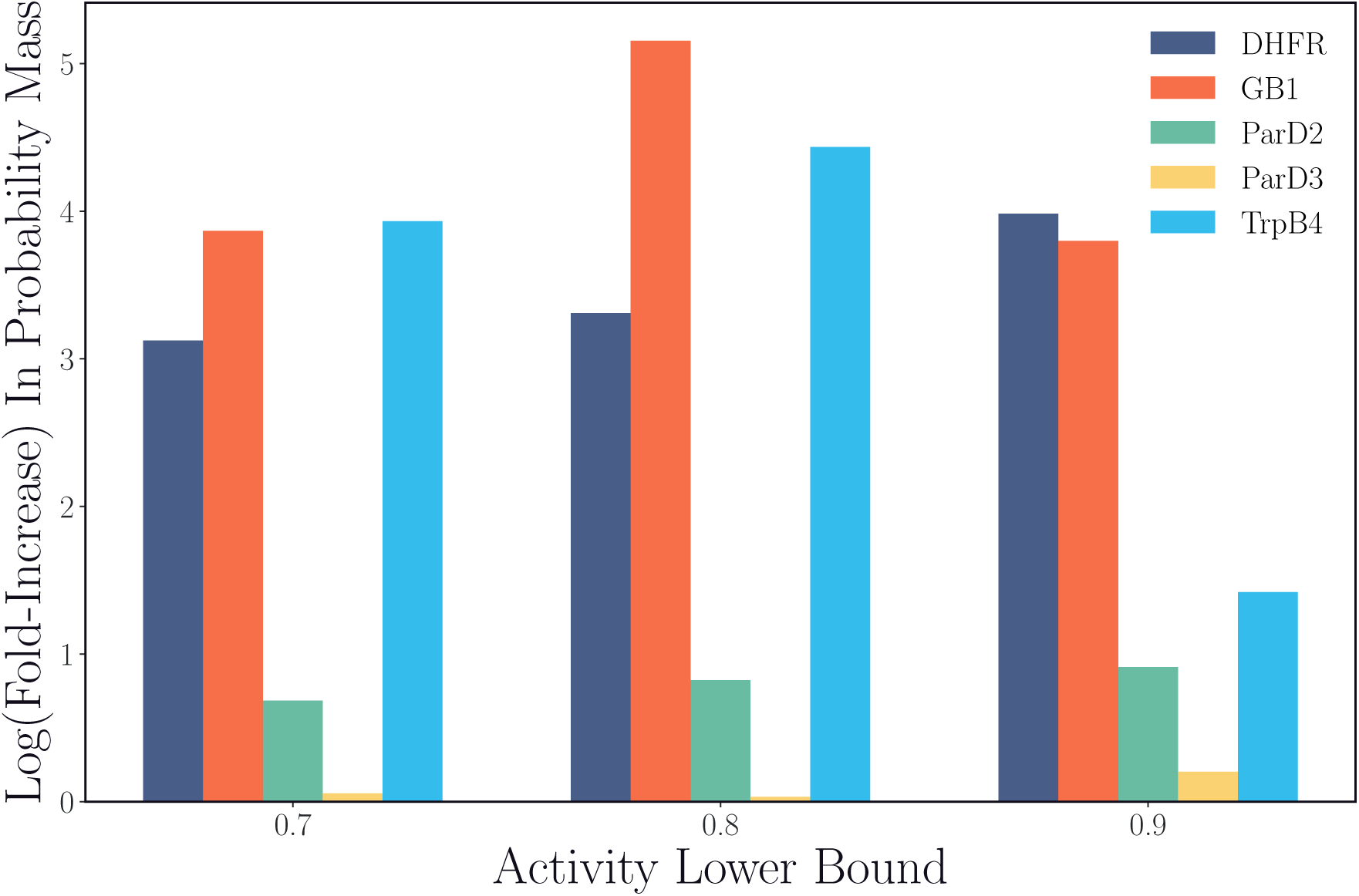
Fold-increase in the probability mass assigned to sequences above a certain experimental activity under *π*_ESM3_ before and after alignment across five multi-site protein fitness landscapes. The post-alignment *π*_ESM3_ is obtained from the final checkpoint of a random iterative alignment experiment.

**Figure 4:**
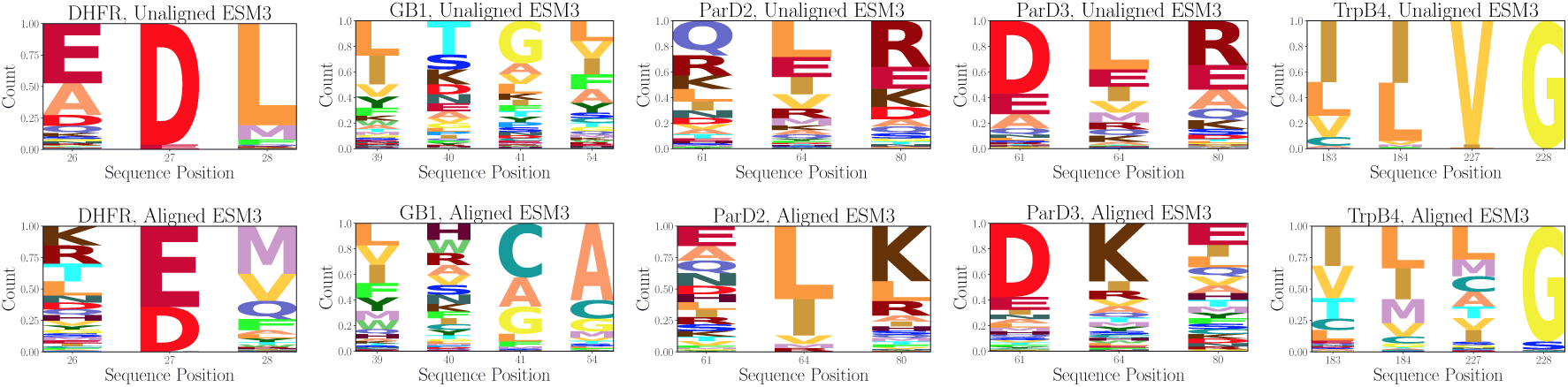
Evolution of multiple sequence alignments constructed using *π*_ESM3_ as relative weights for all possible sequences in five combinatorial multi-site protein fitness landscapes. The top row displays this MSA prior to alignment, and the bottom row shows the MSA after alignment using the final checkpoint from a random iterative alignment experiment.

With the likelihoods from *π*_ESM3_ computed before and after ERA, we use these as relative weights to construct a multiple sequence alignment (MSA) over the landscape of possible mutants at the designated positions for each landscape. Computing the MSA over the aligned and unaligned *π*_ESM3_ shows that the learned representation identifies the residues that are crucial for high experimental activity. For DHFR, ParD2, and ParD3, the most prevalent residues at each position following alignment are the exact residues present at those positions in the optimal mutant. For GB1, the heightened prevalence of cysteine and alanine at positions 41 and 54 reflect the fact that 8 of the 10 most active sequences in the landscape contain alanine at position 54 and either cysteine or alanine at position 41. The other 2 of the top 10 sequences have leucine at position 41 and glycine at position 54, which are also among the most represented residues in the MSA. Moreover, the increased prevalence of tryptophan at position 40 is also promising, given its presence in 7 of the 10 most active sequences in the landscape.

For TrpB4, the preference for I183, L184, and G228 are in line with the optimal sequence in the landscape. The reasons behind these preferences become clearer in the context of the parent sequence for the TrpB4 landscape being an engineered protein itself [22], so these mutations reflect a reversion back toward the naturally occurring *Tm*-TrpB4 protein this was engineered from. However, the optimal mutation to lysine at position 227 is harder to find in the MSA, which we should expect given its less than 1 percent prevalence in the natural MSA for the TrpB protein [22]. Even though lysine is not explicitly favored to the same extent as optimal mutations at other positions, the transition from upweighting valine at position 227 to a less concentrated distribution makes exploration more feasible than with the pre-alignment policy.

These significant distributional shifts are also reflected when sampling from *π*_ESM3_ over multiple rounds of alignment. Figure D.2 demonstrates the percent change in count of residues generated from *π*_ESM3_ from the unaligned ESM3-1.4B policy to the final round of alignment for GB1 and TrpB4. These trends reflect the changes observed in the MSAs. For GB1, there is enhanced prioritization of residues that are crucial for high activity. Though such enhancements are less clear for TrpB4, sampling from *π*_ESM3_ is more exploratory than before ERA.

### 2.5 Additional Conditioning Beyond Primary Sequence

The multi-modal nature of ESM3 enabled us to investigate whether additional conditioning beyond a protein’s primary sequence would help or harm the performance of ERA in optimizing for protein activity. Thermostability is a well-understood prerequisite for any arbitrary protein function, so we aimed to test whether learning more general information about thermostability prior to fine-tuning on a specific experimental dataset would provide ESM3-1.4B with a helpful prior for mutational effects. Connections between sequence likelihoods and thermostability were originally described in [23, 29] and have been recently employed to predict changes in affinity conditioned on structure predictions [9]. To this end, we pre-trained ESM3-1.4B with the full Megascale [43] mutational thermostability dataset using the ERA objective. The dataset contains single and double mutations to over 400 parent sequences, so the number of total comparisons that could be formed was impractical for full fine-tuning. To alleviate this issue, we trained for a single epoch only, and only utilized pairs consisting of one stabilizing and one destabilizing mutation for each parent sequence.

Furthermore, the intrinsic link between 3D protein structure and function suggests that conditioning on backbone coordinates within the ESM3-1.4B model would provide useful information for sequence optimization. The choice of backbone coordinates we made were the ground-truth crystal structures from the PDB [3] for each of the parent sequences in the landscapes of interest, because all the activities being optimized are closely tied to the natural function of each protein. Lastly, though ESM3-1.4B was not trained explicitly with bound complex structures, we also attempted to condition on the backbone coordinates of the bound complex for the three binding affinity landscapes (GB1, ParD2, and ParD3).

Interestingly, neither structure conditioning nor thermostability pre-training substantively improves performance over the pure sequence model. We first examine the change in the log *π*_ESM3_ distributions as a result of the Megascale pretraining and the backbone co-ordinate conditioning. Figure 5 demonstrates that while the overall shape of the log *π*_ESM3_ distributions remain the similar, the spread of log *π*_ESM3_ widens considerably because of this fine-tuning. This is most noticeable in the GB1 landscape, where the least likely sequences become around 30 orders of magnitude less likely than they were previously. Figure 6 demonstrates a shift in probability mass toward higher-activity sequences for GB1, ParD2, and ParD3, while the distribution of probability masses generally does not change much for DHFR and TrpB4. Similar behavior is observed as a result of the backbone coordinate conditioning in Figures 7 and 8. The main difference is that the shift in probability mass for GB1 toward higher-activity sequences is not as pronounced, and that trend is actually reversed for both ParD2 and ParD3.

**Figure 5:**
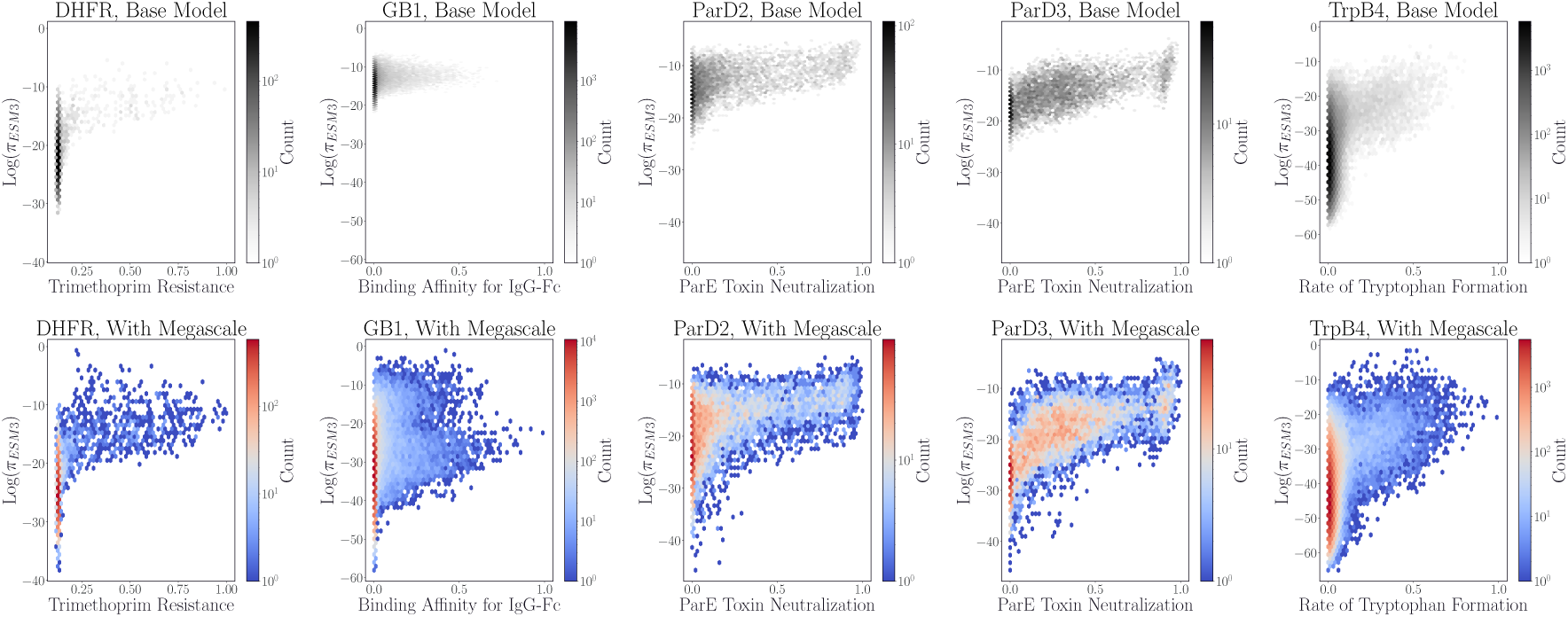
Evolution of the relationship between log *π*_ESM3_ and experimental activity from before (top row) to after (bottom row) ERA pre-training with the Megascale thermostability dataset [43].

**Figure 6:**
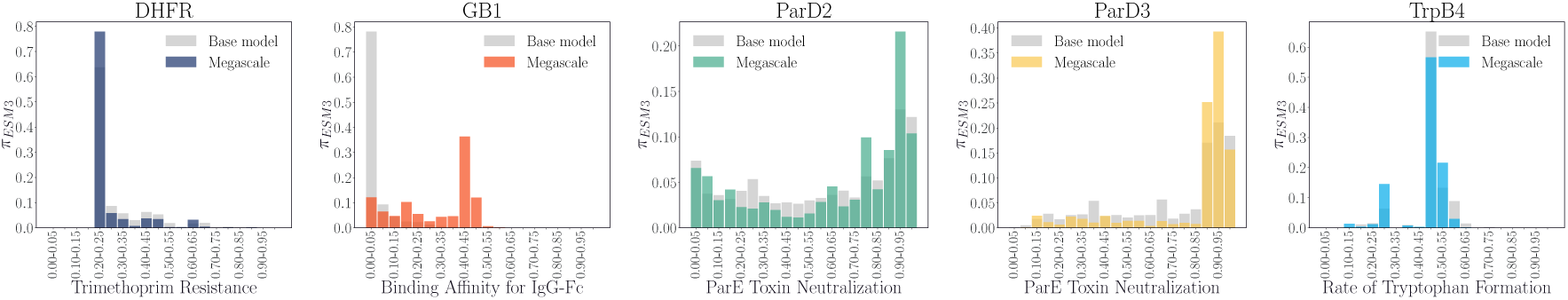
Evolution of probability mass from *π_ESM_*_3_ binned by sequence fitness from before (gray) to after (color) ERA pre-training on the Megascale thermostabilty dataset [43].

**Figure 7:**
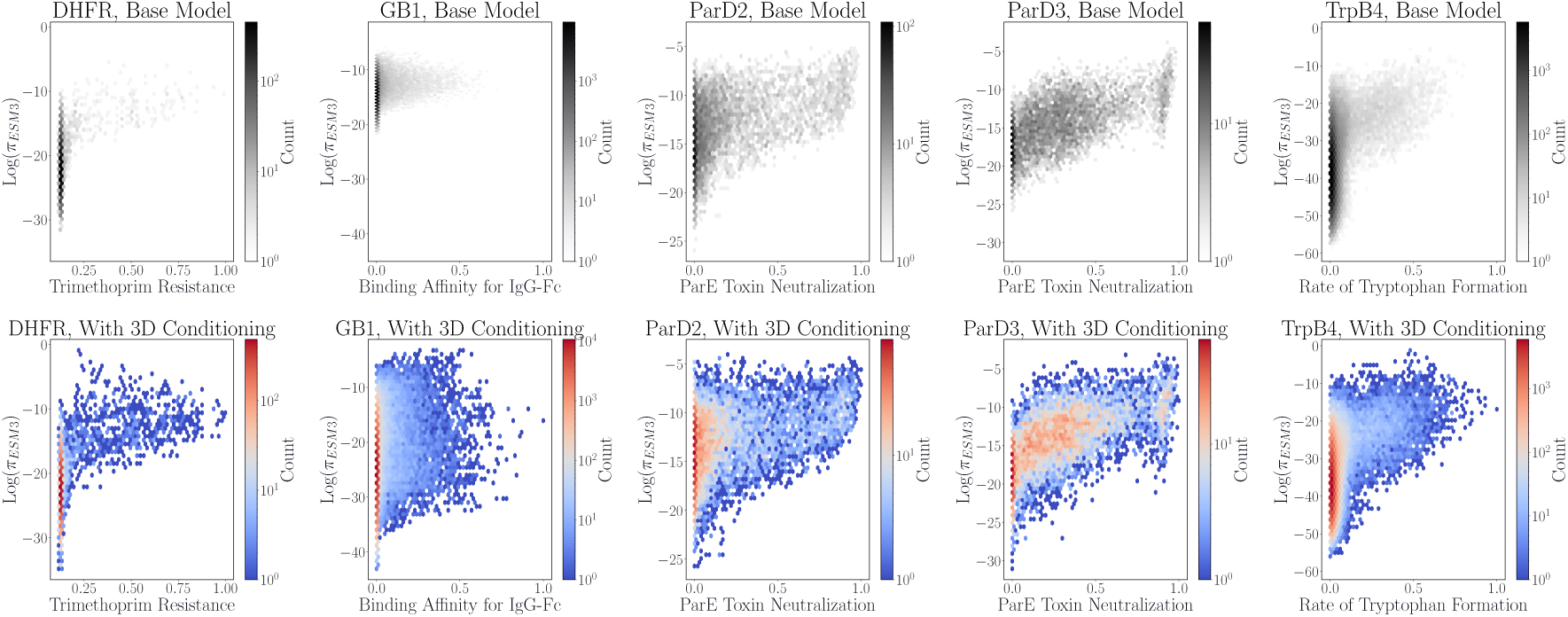
Evolution of the relationship between log *π*_ESM3_ and experimental activity from before (top row) to after (bottom row) conditioning ESM3-1.4B with the backbone 3D coordinates of a protein target’s wild type crystal structure from the PDB.

**Figure 8:**
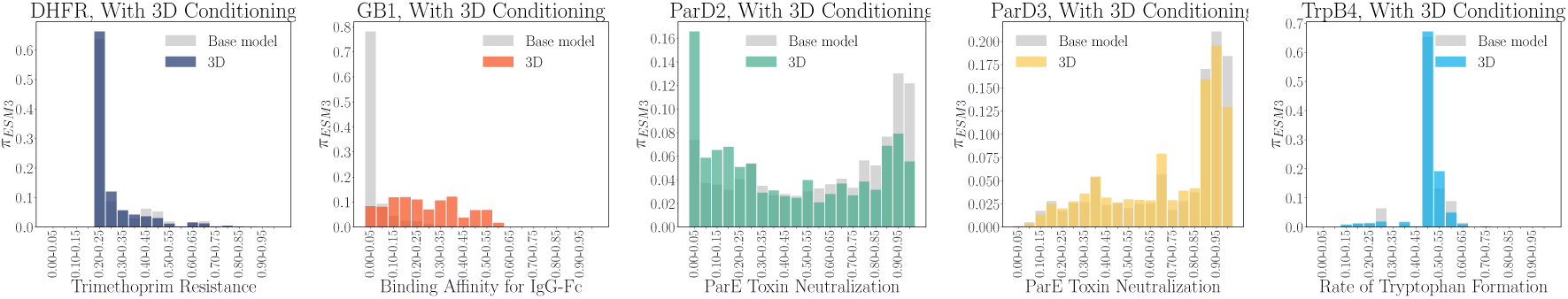
Evolution of probability mass landscapes with conditioning on the backbone coordinates of the wild type protein structure.

Regardless of whether the probability distributions were shifted toward the most active regions of sequence space or not, Tables 7 and 8 suggest that none of these strategies led to an improved ability for ESM3-1.4B to navigate these multi-site, combinatorial landscapes. The shortcomings of the Megascale pretraining strategy can be rationalized by considering that the single- and double-mutant nature of the dataset provides no opportunity to consider higher-order epistatic effects, which are crucial toward effectively navigating rugged landscapes. Regarding the backbone conditioning strategies, those structural constraints are likely too strict for full sequence optimization and make local optima in the fitness landscape harder to escape. This is consistent with previous studies of structure-informed protein language models benefitting less from this conditioning on near-native optimization tasks [34], as in the five landscapes we explore here.

**Table 7:** Average maximum fitness achieved across multiple replicate runs with standard errors included for different variations of ERA on ESM3-1.4B. In addition to testing ERA with only the sequence track of ESM3-1.4B, we also attempted conditioning on the 3D coordinates of the wild type backbone, the 3D coordinates of the bound complex (for binding landscapes), and pretraining with ERA on the Megascale thermostability dataset [43]. All methods used 384 total samples over four rounds for training, with the results being averaged over 10 replicates.

| Method | DHFR | GB1 | ParD2 | ParD3 | TrpB4 |
| --- | --- | --- | --- | --- | --- |
| ERA | **1.0 ± 0.00** | **0.83 ± 0.04** | **1.0 ± 0.00** | **1.0 ± 0.00** | **0.91 ± 0.02** |
| ERA (w/ struct. cond.) | - | 0.76 ± 0.04 | - | - | 0.89 ± 0.02 |
| ERA (w/ complex) | - | 0.74 ± 0.04 | **1.0 ± 0.00** | 0.99 ± 0.00 | - |
| ERA (w/ stab. pretrain.) | **1.0 ± 0.00** | 0.78 ± 0.04 | **1.0 ± 0.00** | 0.99 ± 0.00 | 0.85 ± 0.02 |

**Table 8:** Fraction of replicate runs reaching the maximum fitness sequence in the landscape for different variations of ERA on ESM3-1.4B. In addition to testing ERA with only the sequence track of ESM3-1.4B, we also attempted conditioning on the 3D coordinates of the wild type backbone, the 3D coordinates of the bound complex (for binding landscapes), and pretraining with ERA on the Megascale thermostability dataset [43]. All methods used 384 total samples over four rounds for training, with the results being averaged over 10 replicates.

| Method | DHFR | GB1 | ParD2 | ParD3 | TrpB4 |
| --- | --- | --- | --- | --- | --- |
| ERA | **1.0** | **0.20** | **1.0** | **0.90** | **0.30** |
| ERA (w/ struct. cond.) | - | 0.10 | - | - | 0.10 |
| ERA (w/ complex) | - | 0.10 | **1.0** | 0.70 | - |
| ERA (w/ stab. pretrain.) | - | **0.20** | **1.0** | 0.50 | 0.10 |

Despite these modifications to the method being ineffective, there are other avenues to improving performance that leverage information learned by the model to make sampling more efficient. For instance, one can identify motifs that are over-represented in the most active sequences empirically, such as in Figures D.3-D.7. Sampling can be conducted with certain clearly advantageous residues fixed, or restricted to a certain Hamming distance away from top sequences in previous rounds. We have found this to be an effective strategy for improving full sequence optimization, leading to 6 out of 10 replicates reaching the optimal TrpB4 sequence. The ability to examine past generations coupled with the learned distribution of the aligned *π*_ESM3_ offers an avenue through which more informed decisions about sampling can be made.

## 3 Conclusions

As powerful foundation models emerge across scientific disciplines, it is imperative that we understand what these representations offer for generalization from sparse data. Sequence-based representations of proteins are obviously quite limited, lacking direct structural information, detailed chemical and physical priors, or explicit constraints on thermostability. Nevertheless, we find here that these representations provide a robust starting point for design tasks, even when starting from an engineered protein. Our approach, which leverages experimental data to shift the sequence generation probabilities, learns on-the-fly to adapt the latent information in a protein language model towards desired function.

How to construct optimal representations for experimentally driven design tasks is a major question looking forward. While our experiments suggest that incorporating additional non-sequence conditioning (e.g., providing 3D coordinates, or pre-training on thermostability data) does not systematically improve predictions, increasingly rich representations may help to overcome thorny issues like epistasis. Indeed, incorporating the inductive bias of a protein language model, which selects sequences based on evolutionary likelihood, does not provide a strong prior for arbitrary protein optimization tasks. A more comprehensive assessment of domain generalization of our approach would help determine if this additional information would benefit design in contexts constrained by structure.

Finally, while the expressive large-scale neural networks we use in this study yield state-of-the-art results, the computational costs of fine-tuning these models are high in comparison to the minimal architectures used by other ML-driven DE approaches. Our focus was to determine the limits of the approach and hence we did not explore parameter efficient fine-tuning methods. These approaches are increasingly understood to yield results on par with full fine-tuning but at substantially lower costs, a topic we will explore in future work.

## Acknowledgements

SI acknowledges support from the National Science Foundation Graduate Research Fellowship Program and the Shoucheng Zhang Graduate Fellowship. Research reported in this publication was supported by the National Institute of General Medical Sciences of the National Institutes of Health under award number 1R35GM159834-01. The content is solely the responsibility of the authors and does not necessarily represent the official views of the National Institutes of Health.

## Data and Code Availability

The code used for training, sampling, and likelihood calculations is available at GitHub (rotskoff-group/era-directed-evolution). Model checkpoints and iterative alignment results are on Hugging Face: dataset and models.

## A Training Hyperparameters

In accordance with preliminary results [6, 7], ERA hyperparameters of *β* = 10.0 and *γ* = 0 were used for all experiments. We found further increases to *β* did not provide appreciable benefits to performance, and increases to *γ* were detrimental. For the experiments here, we used the RMSProp optimizer with a learning rate of 1.0 ∗ 10^−5^ and trained for 25 epochs. The only exception was the Megascale pretraining experiment, which was only trained for a single epoch given the amount of preference pairs present in the overall dataset.

Given the relationship between *β*_DPO_ and *β*_ERA_ established in [7], we ran all DPO experiments with *β*_DPO_ = 1.0. Like with the ERA experiments, we used the RMSProp optimizer with a learning rate of 1.0 ∗ 10^−5^ and trained for 25 epochs.

The random forest hyperparameters and sequence-pooling of ESM2-650M embeddings described in [20] were followed exactly in the benchmark of the method performed in this study. Training was performed on a single Nvidia A100 16GB GPU.

For all ESM3-1.4B experiments, we used resources of the National Energy Research Scientific Computing Center (NERSC), a Department of Energy Office of Science User Facility. Jobs run on NERSC used 4 Nvidia A100 GPUs (either 40GB or 80GB depending on what was allocated).

## B Conditional Sequence Probability Distribution Metrics

For a fixed parent sequence and a fixed set of masked positions, we sample *N* mutated sequences *x*_1_, *x*_2_, …, *x_N_* and compute their joint *π*_ESM3_ according to 1. We define the normalized sequence probabilities as

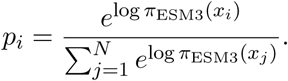

### Sequence entropy

The entropy of the induced sequence distribution is

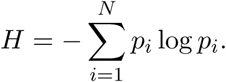

Lower entropy indicates a more concentrated, mode-seeking distribution over sequences.

### KL divergence to a uniform distribution

To quantify deviation from a uniform distribution over the sampled sequences, we compute

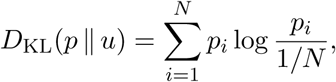

where *u_i_* = 1*/N*. Larger values indicate a more peaked, non-uniform distribution.

### Top-*k* cumulative probability mass

We also measure the fraction of total probability mass captured by the *k* most likely sequences. Let *p*_(1)_ ≥ *p*_(2)_ ≥ · · · ≥ *p*_(*N*)_ denote the probabilities sorted in descending order. The top-*k* coverage is defined as

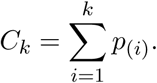

Higher values of *C_k_* indicate that probability mass is concentrated on a small number of sequences.

### Sampling redundancy

Given *N* sampled sequences and *N*_unique_ unique sequences among them, we define the redundancy rate as

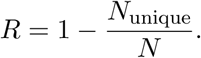

Higher redundancy reflects repeated sampling of the same high-probability sequences and is indicative of mode-seeking behavior.

## C PDB Identifiers for Backbone Structures Used As Conditioning

| Landscape | PDB Identifier |
| --- | --- |
| DHFR | 6XG5 |
| GB1 | 2GI9 |
| ParD2 | 6X0A |
| ParD3 | 5CEG |
| TrpB4 | 8VHH |

## D Visualization of Residue Sampling Frequencies

**Figure D.1:**
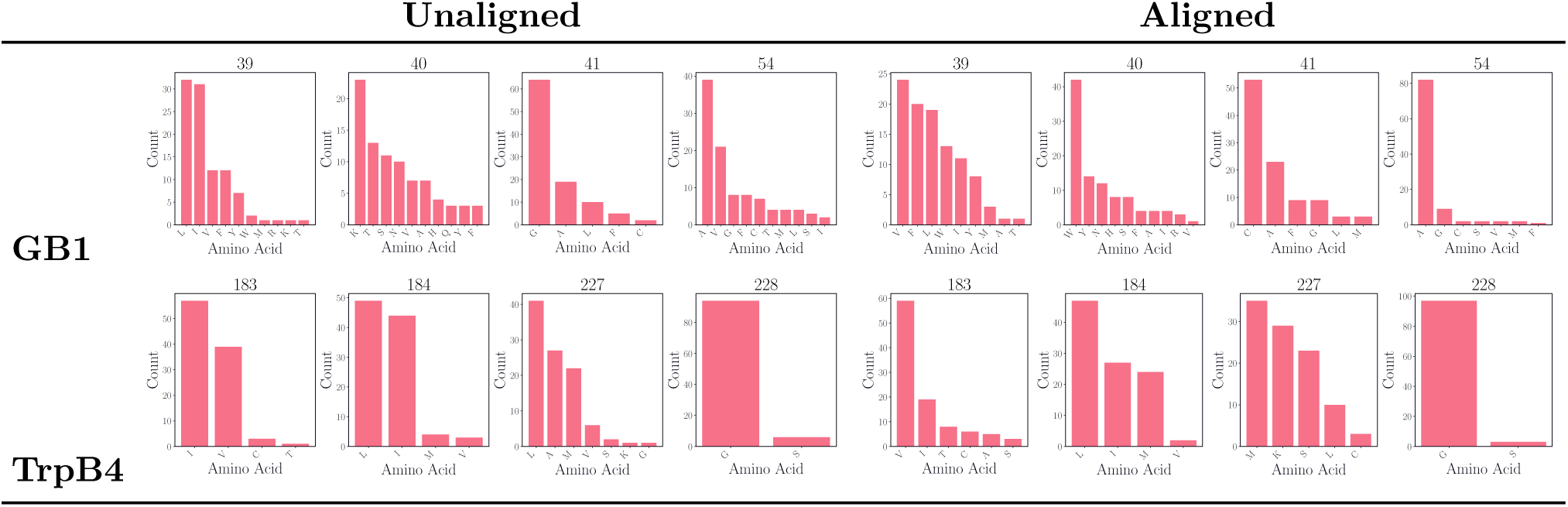
Frequency of occurrence of amino acid residues in the four-site GB1 and TrpB4 landscapes among the top 100 sequences by experimental fitness over 10 replicates of 96 samples. The plots on the left half are obtained when sampling from the unaligned *π*_ESM3_, while the plots on the right half are sampled from the final checkpoints after four rounds of alignment with 96 new sequences for training at each round.

**Figure D.2:**
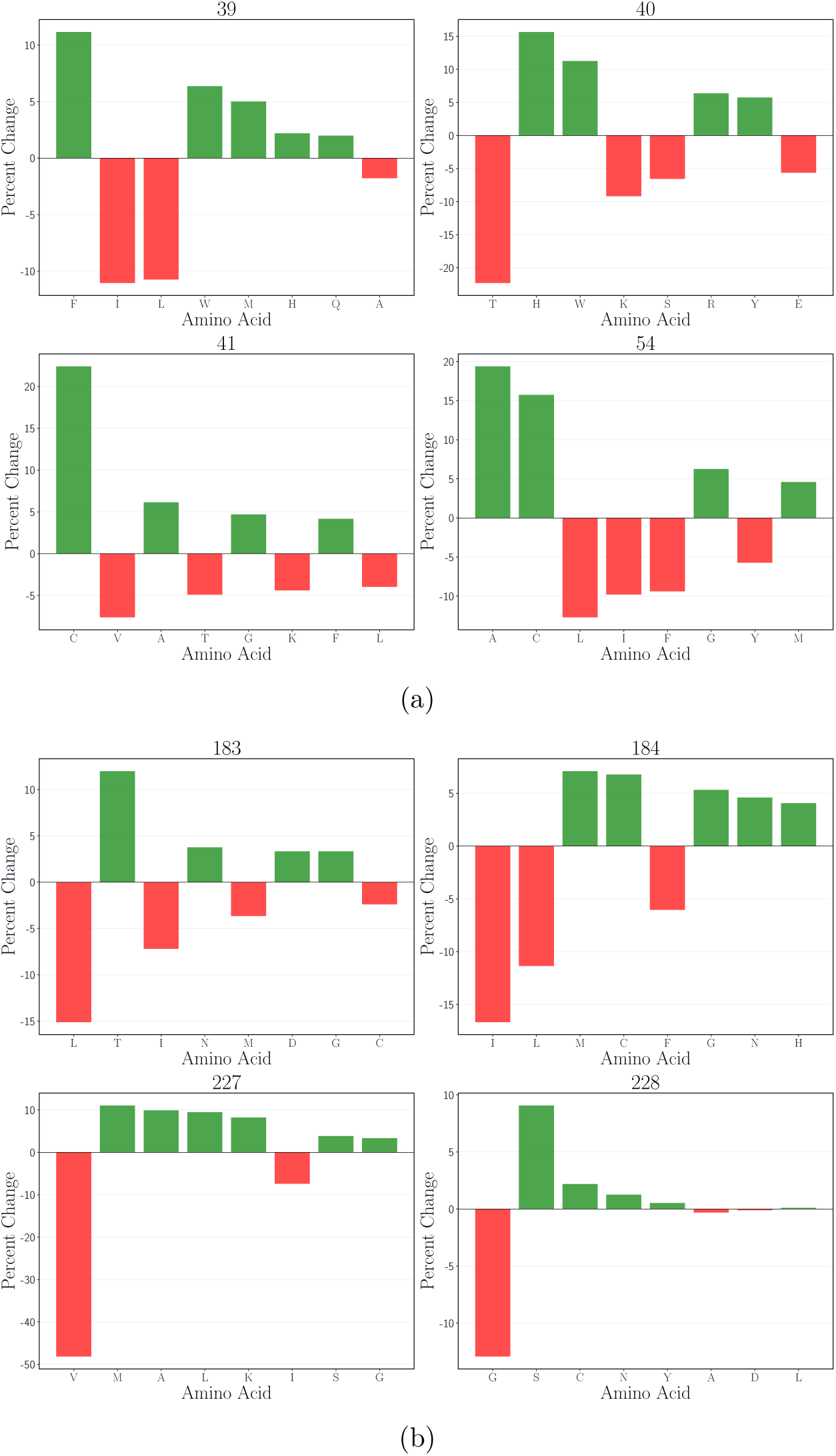
Change in the frequencies of amino acid residues in 96 generated sequences from the unaligned *π*_ESM3_ to the aligned *π*_ESM3_ after four rounds of alignment with 96 sequences each. The top and bottom panels display the discrepancies for the GB1 and TrpB4 experiments, respectively. The discrepancies are computed across 10 replicate runs of the iterative alignment experiments, where no duplicate or previously trained sequences are permitted during sampling.

**Figure D.3:**
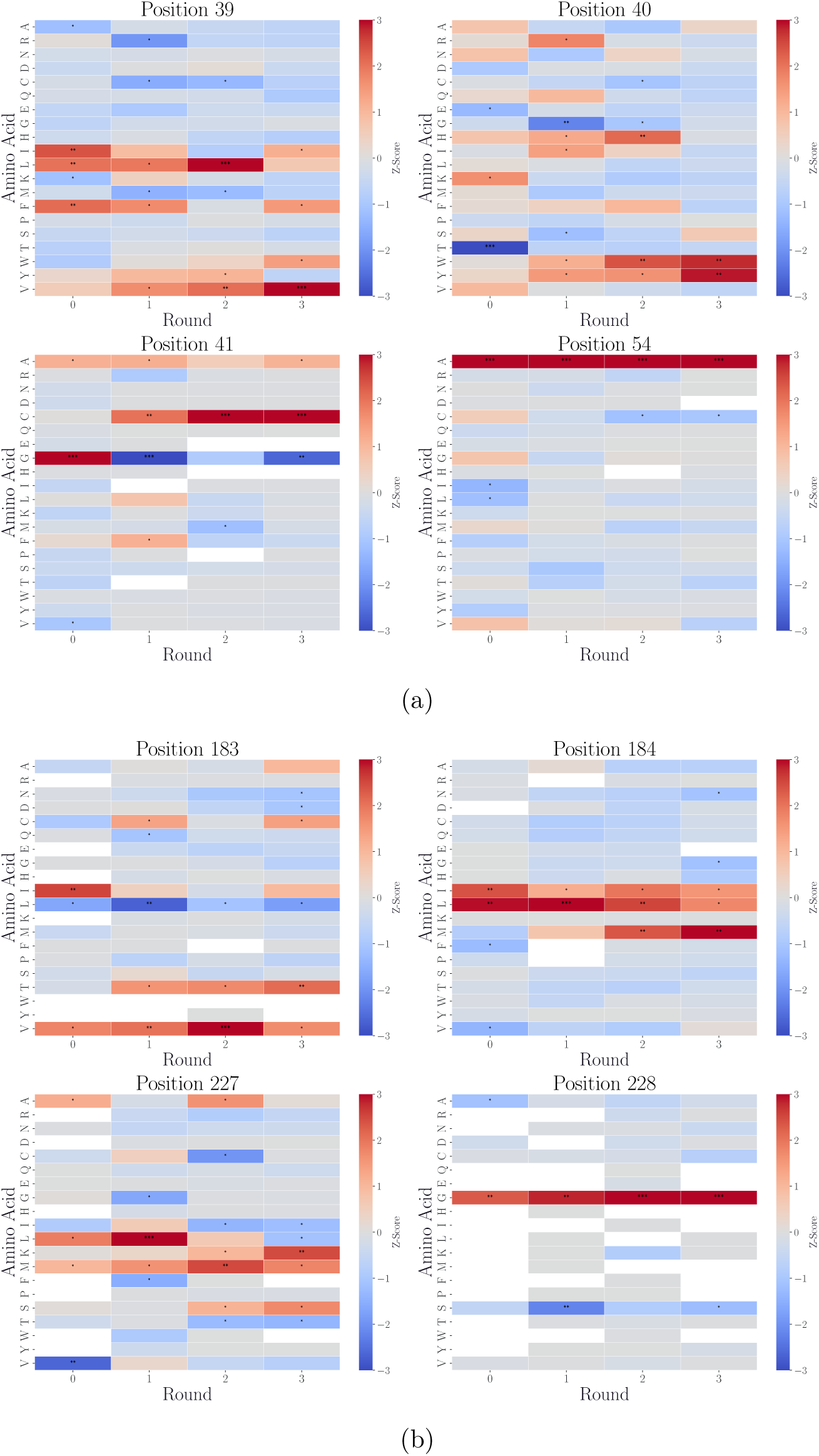
Heatmap of Z-scores of the discrepancy between the occurrence of residues in the top 100 sequences by fitness compared to all other sequences sampled at each round of alignment over 10 replicate iterative alignment experiments. The top and bottom panels visualize the Z-scores for the GB1 and TrpB4 experiments, respectively.

**Figure D.4:**
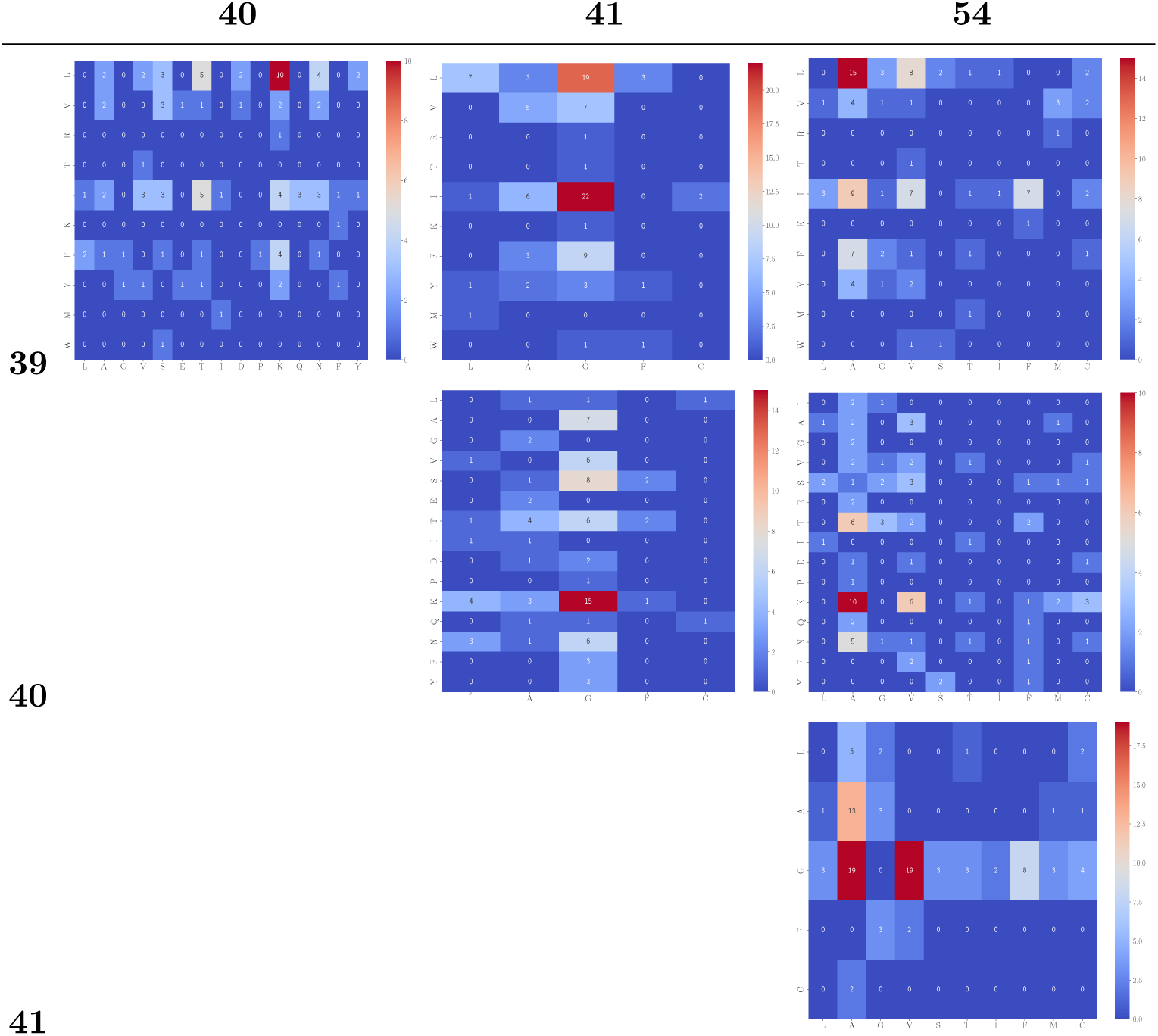
Co-occurrence of pairs of residues from top 100 sequences by experimental fitness over 10 replicates of GB1 sampling from unaligned *π*_ESM3_. A total of 96 sequences were sampled in each replicate. Rows and columns correspond to residue positions (39, 40, 41, 54). Only the upper triangle is shown because co-occurrence is symmetric with respect to position order.

**Figure D.5:**
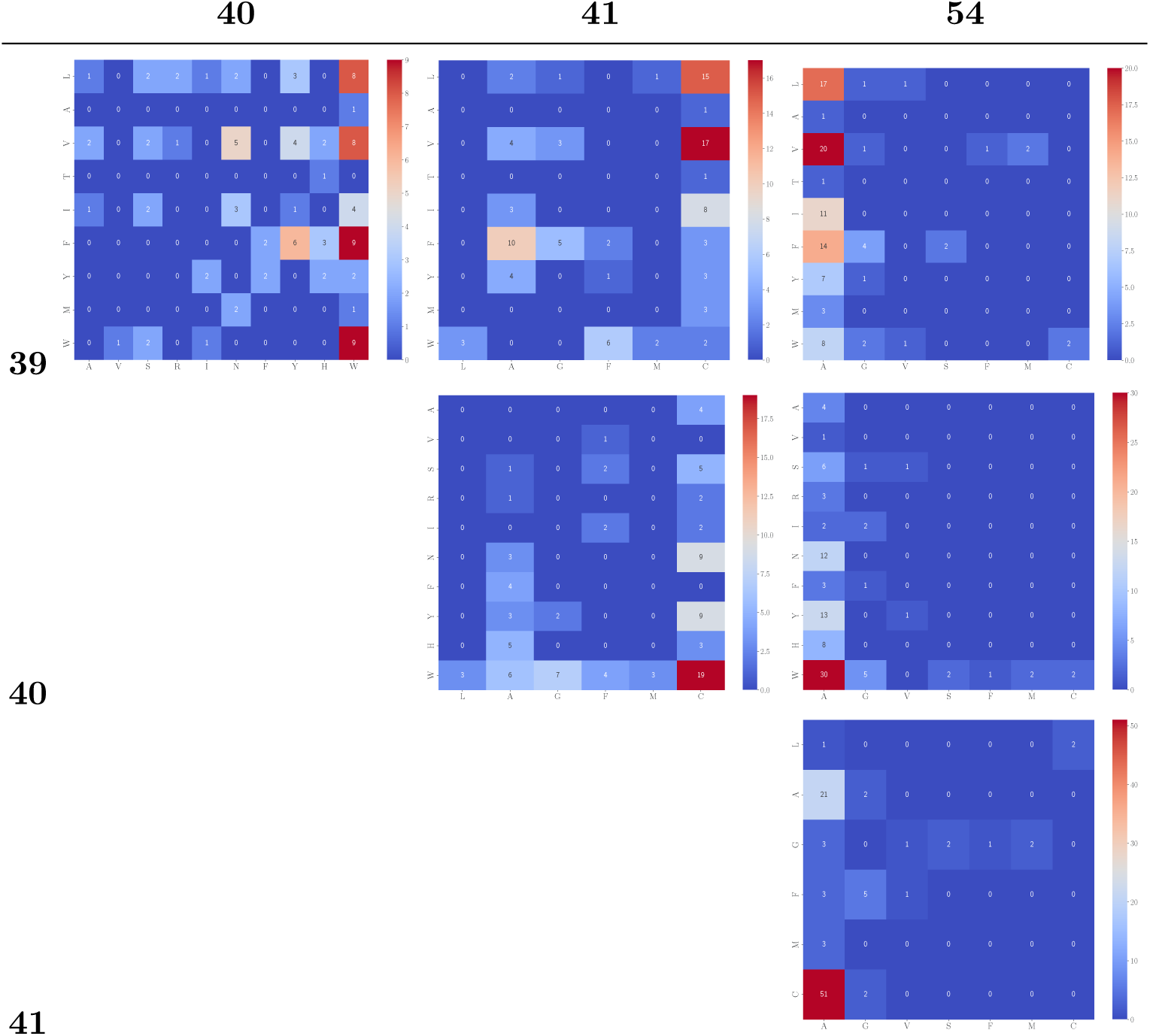
Co-occurrence of pairs of residues from top 100 sequences by experimental fitness over 10 replicates of GB1 sampling from a *π*_ESM3_ aligned with 384 sequences over 4 rounds. A total of 96 sequences were sampled in each replicate. Rows and columns correspond to residue positions (39, 40, 41, 54). Only the upper triangle is shown because co-occurrence is symmetric with respect to position order.

**Figure D.6:**
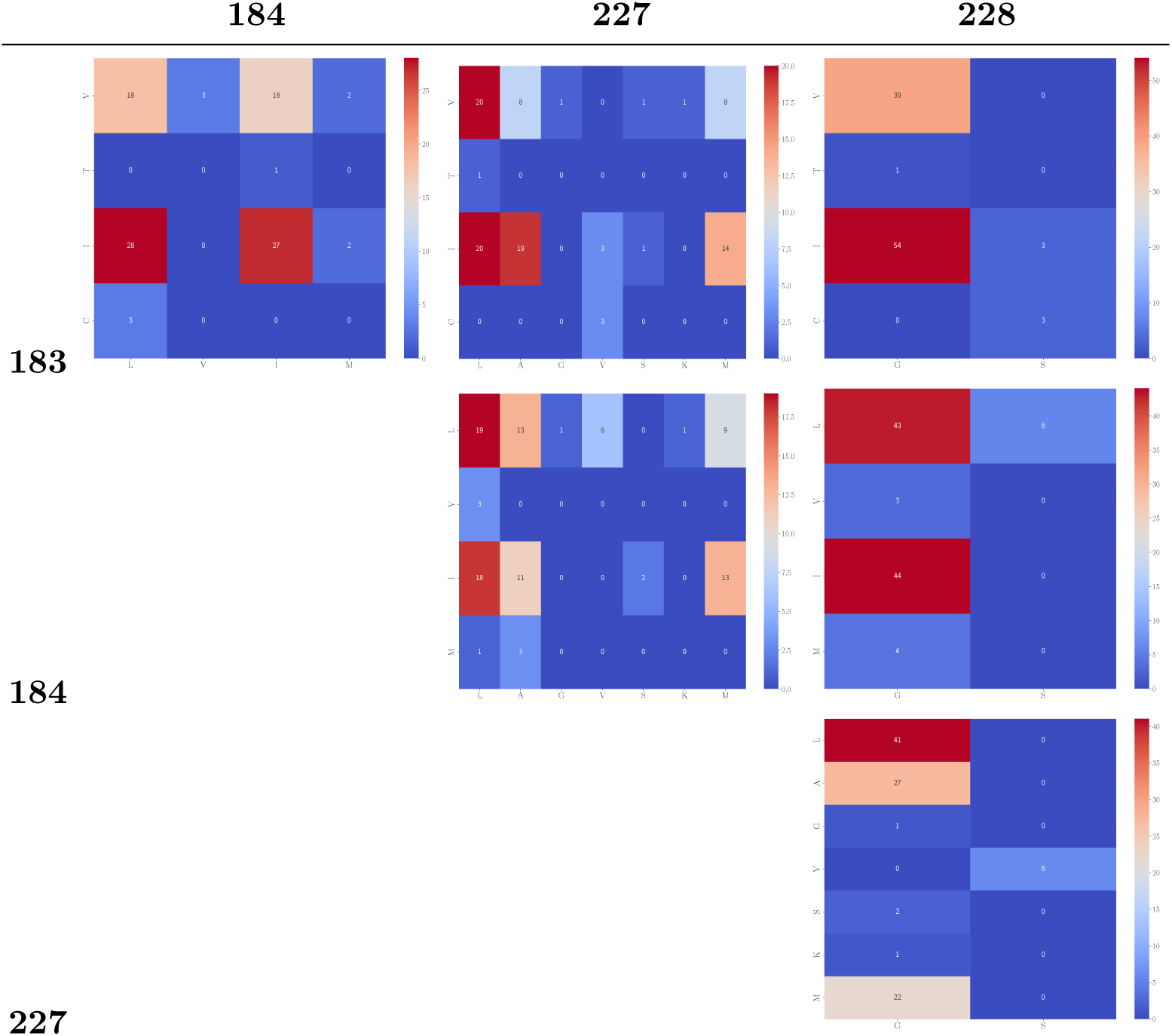
Co-occurrence of pairs of residues from top 100 sequences by experimental fitness over 10 replicates of TrpB4 sampling from unaligned *π*_ESM3_. A total of 96 sequences were sampled in each replicate. Rows and columns correspond to residue positions (183, 184, 227, 228). Only the upper triangle is shown because co-occurrence is symmetric with respect to position order.

**Figure D.7:**
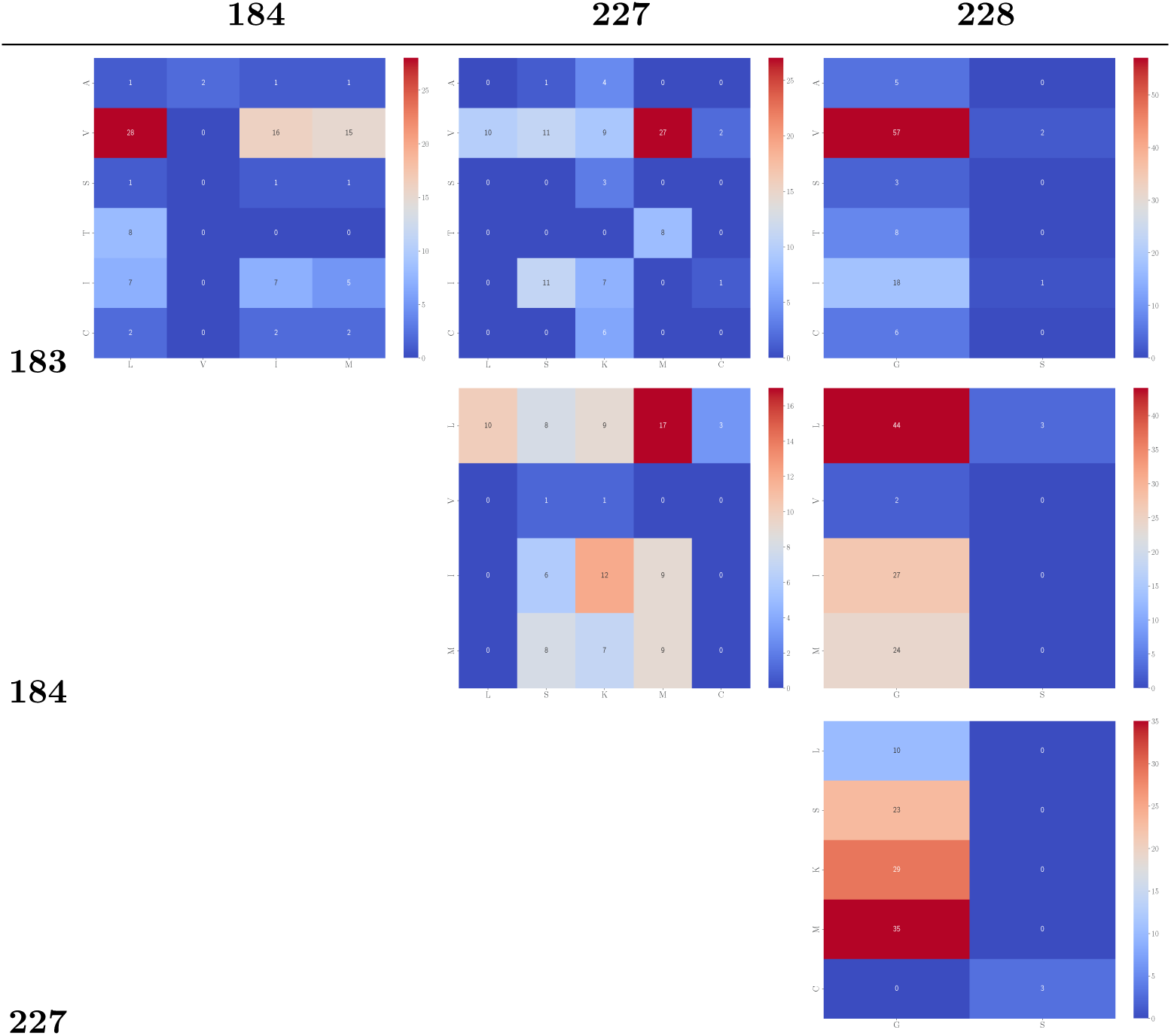
Co-occurrence of pairs of residues from top 100 sequences by experimental fitness over 10 replicates of TrpB4 sampling from a *π*_ESM3_ aligned with 384 sequences over 4 rounds. A total of 96 sequences were sampled in each replicate. Rows and columns correspond to residue positions (183, 184, 227, 228). Only the upper triangle is shown because co-occurrence is symmetric with respect to position order.

